# Disrupted gene networks in subfertile hybrid house mice

**DOI:** 10.1101/776286

**Authors:** Katy Morgan, Bettina Harr, Michael A. White, Bret A. Payseur, Leslie M. Turner

## Abstract

The Dobzhansky-Muller model provides a widely accepted mechanism for the evolution of reproductive isolation: incompatible substitutions disrupt interactions between genes. To date, few candidate incompatibility genes have been identified, leaving the genes driving speciation mostly uncharacterized. The importance of interactions in the Dobzhansky-Muller model suggests that gene coexpression networks provide a powerful framework to understand disrupted pathways associated with postzygotic isolation. Here, we perform Weighted Gene Coexpression Network Analysis (WGCNA) to infer gene interactions in hybrids of two recently diverged European house mouse subspecies, *Mus mus domesticus* and *M. m. musculus*, which commonly show hybrid male sterility or subfertility. We use genome-wide testis expression data from 467 hybrid mice from two mapping populations: F_2_s from a laboratory cross between wild-derived pure subspecies strains and offspring of natural hybrids captured in the Central Europe hybrid zone. This large data set enabled us to build a robust consensus network using hybrid males with fertile phenotypes. We identify several expression modules, or groups of coexpressed genes, that are disrupted in subfertile hybrids, including modules functionally enriched for spermatogenesis, cilium and sperm flagellum organization, chromosome organization and DNA repair, and including genes expressed in spermatogonia, spermatocytes and spermatids. Our network-based approach enabled us to hone in on specific hub genes likely to be influencing module-wide gene expression and hence potentially driving Dobzhansky-Muller incompatibilities. A total of 69 (24.6%) of these genes lie in sterility loci identified previously in these mapping populations, and represent promising candidate barrier genes and targets for future functional analysis.

## Introduction

According to the classic Dobzhansky-Muller (DM) model of speciation, mutations that accumulate independently and in different genomic regions may be incompatible when brought together in a hybrid background, resulting in disrupted epistasis and the development of postzygotic reproductive barriers (Dobzhansky 1982; Muller, 1942). These barriers, which include reductions in hybrid fertility and/or hybrid viability, in turn restrict gene flow, enabling the further divergence of incipient species. The applicability of the DM model to allopatric speciation scenarios has been demonstrated through both theoretical and empirical studies (reviewed in Presgraves, 2010). However, while many genes and loci influencing hybrid fertility have been described (reviewed in Maheshwari and Barbash, 2011), characterizing the specific interactions between loci underpinning Dobzhansky-Muller incompatibilities (DMIs) remains a challenge.

While DMIs were originally assumed to act independently of one another, such that individual loci are involved in a single DMI (Orr, 1995), accumulating evidence raises questions about this assumption. A computational modeling study based on RNA folding demonstrated that as two populations evolve in allopatry, DMIs become increasingly complex over time (Kalirad and Azevedo, 2017). In this context, the majority of DMIs could involve more than two loci and individual loci are expected to participate in multiple DMIs (Kalirad and Azevedo, 2017). Evidence from empirical studies in house mice (Turner *et al*. 2014; Turner and Harr 2014), *Drosophila* (Phadnis *et al*. 2011), as well as plant and fungal taxa, supports the prevalence of complex DMIs involving multiple partners (reviewed in Fraïse *et al*. 2014). Theoretical studies have also implicated the role of divergence in complex regulatory pathways in driving postzygotic reproductive isolation via DMIs, with reproductive isolation being more likely to develop when regulatory pathways contain larger numbers of loci (Johnson and Porter, 2000) and are under the influence of directional selection (Porter and Johnson, 2002; Johnson and Porter, 2007). Empirical evidence from house mice suggests that divergence in regulatory elements between subspecies disrupts epistatic interactions between sets of genes, resulting in significant reductions or elevations of gene expression in hybrid relative to pure subspecies individuals (Mack *et al*. 2016).

Misexpression in hybrid individuals is commonly observed, and has been associated with reduced fertility in house mice (Good *et al*. 2010; Turner *et al*. 2014; Turner and Harr 2014; Mack *et al*. 2016; Larson *et al*. 2017), *Drosophila* (Michalak and Noor, 2003; Moehring *et al*. 2006; Gomes and Civetta, 2015), and cats (Davis *et al*. 2015). Understanding the role of potentially complex DMIs in driving patterns of misexpression may be facilitated by exploring gene interactions in a network context, in which expression patterns of sets of genes are allowed to depend upon one another. Network-based approaches cluster genes into coexpression modules, which are likely to be associated with common biological pathways and functions (e.g. Ayroles *et al*. 2009; Miller *et al*. 2010; reviewed in Mackay, 2014), and have been employed to identify sets of genes associated with fitness-related phenotypes (e.g. Jumbo-Lucioni *et al*. 2010; Mack *et al*. 2018) and genes with disrupted interactions associated with disease (e.g. Saris *et al*. 2009; Miller *et al*. 2010; reviewed in de la Fuente, 2010). Identifying the most highly-connected “hub genes” within modules can facilitate identification of changes driving disruption of pathways from downstream misexpressed genes. Network approaches have also been used to identify mechanisms involved in adaptive ecological divergence (e.g. Kelley *et al*. 2016; Gould *et al*. 2018; Zhao *et al*. 2019), and to explore the mechanisms underlying ecological speciation of sympatric lake whitefish ecotypes (Filteau *et al*. 2013). However, the power of network-based analyses for exploring the disrupted gene interactions associated with reduced hybrid fertility and postzygotic reproductive isolation is yet to be realized.

Because mutations and incompatibilities continue to accumulate between independently evolving lineages over time, identifying the incompatibilities that initially caused reproductive isolation requires studying recently diverged lineages, with incomplete reproductive barriers. The European house mouse (*Mus musculus*) provides one such system. Three subspecies, *M. m. musculus, M. m. domesticus* and *M. m. castaneus*, diverged approximately 500,000 years ago (Geraldes *et al*. 2011). Following this initial divergence, *M. m. musculus* and *M. m. domesticus* (henceforth referred to as *musculus* and *domesticus*, respectively) used different routes to colonize Europe, providing the opportunity for the accumulation of DMIs in allopatry. Laboratory crosses between *musculus* and *domesticus* have demonstrated reduced fertility in hybrid males (Forejt and Iványi, 1974; Oka *et al*. 2004; Good *et al*. 2008a), although the degree of sterility varies depending on the nature of the cross (Britton-Davidian *et al*. 2005; Vyskočilová *et al*. 2005; Good *et al*. 2008b; Larson *et al*. 2018). Genetic studies of *musculus-domesticus* hybrids have also led to the first characterization of a mammalian hybrid incompatibility gene, *Prdm9* (Mihola *et al*. 2009), an autosomal histone methyltransferase which binds DNA at recombination hotspots and plays an important role in the initiation of meiotic recombination (Baudat et al. 2010; Davies *et al*. 2016). Negative interactions between some *domesticus Prdm9* alleles and loci on the *musculus* X-chromosome disrupt expression of the X-chromosome and chromosome synapsis, resulting in meiotic arrest (Bhattacharyya *et al*. 2013; Campbell *et al*. 2013; Bhattacharyya *et al*. 2014; Larson *et al*. 2017).

Following colonization of Europe, *musculus* and *domesticus* expanded their ranges to form a zone of secondary contact traversing Central Europe (Macholan *et al*. 2003; Janoušek *et al*. 2012). Hybridization along this secondary contact zone is common, resulting in a cline of genomic admixture (Payseur *et al*. 2004; Macholan *et al*. 2007; Teeter *et al*. 2008; Janoušek *et al*. 2012). Reduced male fertility is frequent in wild-caught hybrids, but complete sterility is rare (Britton-Davidian *et al*. 2005; Albrechtová *et al*. 2012; Turner *et al*. 2012), suggesting that this postzygotic barrier reduces gene flow and contributes to maintenance of the subspecies boundary, but is incomplete and variable in strength across populations.

In the present study, we are building on genetic mapping of fertility phenotypes & gene expression traits in F_2_ *musculus-domesticus* hybrids generated through a laboratory cross (White et al. 2011; Turner *et al*. 2014), and in offspring of wild-caught *musculus-domesticus* hybrids (Turner and Harr 2014), which have identified many autosomal and X-linked sterility loci. DMIs were mapped in both studies, using different methods, yet many loci and interactions are shared between mapping populations (Turner and Harr 2014). Most loci have multiple interaction partners, supporting the presence of complex DMIs (Turner and Harr 2014; Turner *et al*. 2014). Identifying the underlying mechanisms and causative genes remains a challenge, because many loci encompass a large number of genes.

Here, we characterize disruptions in gene networks associated with hybrid male sterility in mice, by analyzing patterns of gene coexpression using microarray data from a total of 467 mice from two *musculus-domesticus* hybrid mapping populations (Turner and Harr 2014; Turner *et al*. 2014). We use weighted gene coexpression analysis (WGCNA, Langfelder and Horvarth, 2008) to (1) characterize a consensus ‘fertile’ network, (2) identify modules of coexpressed genes associated with biological pathways and processes, (3) identify modules disrupted in subfertile hybrids, and (4) identify specific candidate genes likely to be driving network disruptions in subfertile hybrids.

## Results

### Concordant genome-wide testis expression patterns in F_2_ and HZ hybrids

We analyzed genome-wide expression patterns in testes from 295 F_2_ hybrids between two inbred strains of *musculus* and *domesticus* (White et al. 2011; Turner *et al*. 2014) and 172 lab-bred male offspring of mice wild-caught from a hybrid zone (Turner and Harr 2014). Principal components analysis (PCA) of the F_2_ and Hybrid Zone (HZ) data sets showed similar overall patterns (Supplementary Figure 1). For both populations, PC1, which explains 20.1% of variation in F_2_ hybrids and 27.8% of variation in HZ hybrids, is clearly associated with fertility classified on the basis of relative testis weight (testis weight/body weight) and sperm count (White et al. 2011; Turner et al. 2012; Supplementary Figure 1A-B). The PC1 loadings for 25,146 probes in the two datasets are strikingly highly correlated (r=0.91, p<2.2x10^-16^), suggesting a common set of genes contribute to the “sterile” expression pattern (Supplementary Figure 1C). The remarkable similarities in overall testis expression pattern between mapping populations, and previous evidence for shared incompatibilities (Turner and Harr 2014), motivated us to combine expression data from F_2_ and HZ hybrids to characterize gene network disruptions associated with sterility.

After combining and batch-correcting data from F_2_ and HZ hybrid and pure subspecies males, PC1 explains 18.26% of variation in gene expression and again is clearly associated with fertility (Figure 1). The PC1 scores of fertile hybrid and pure subspecies males show similar levels of variation (variance: 609.36 and 616.10, respectively), while variation in PC1 scores is considerably higher in hybrids with subfertile phenotypes (variance: 15149.70). Hybrids for which both fertility phenotypes fall within the pure subspecies range, yet at least one of the phenotypes is more than one standard deviation from the pure subspecies mean, were classified as “intermediate phenotype (see Methods) and show intermediate levels of variation along PC1 (variance: 1816.78).

**Fig 1.**
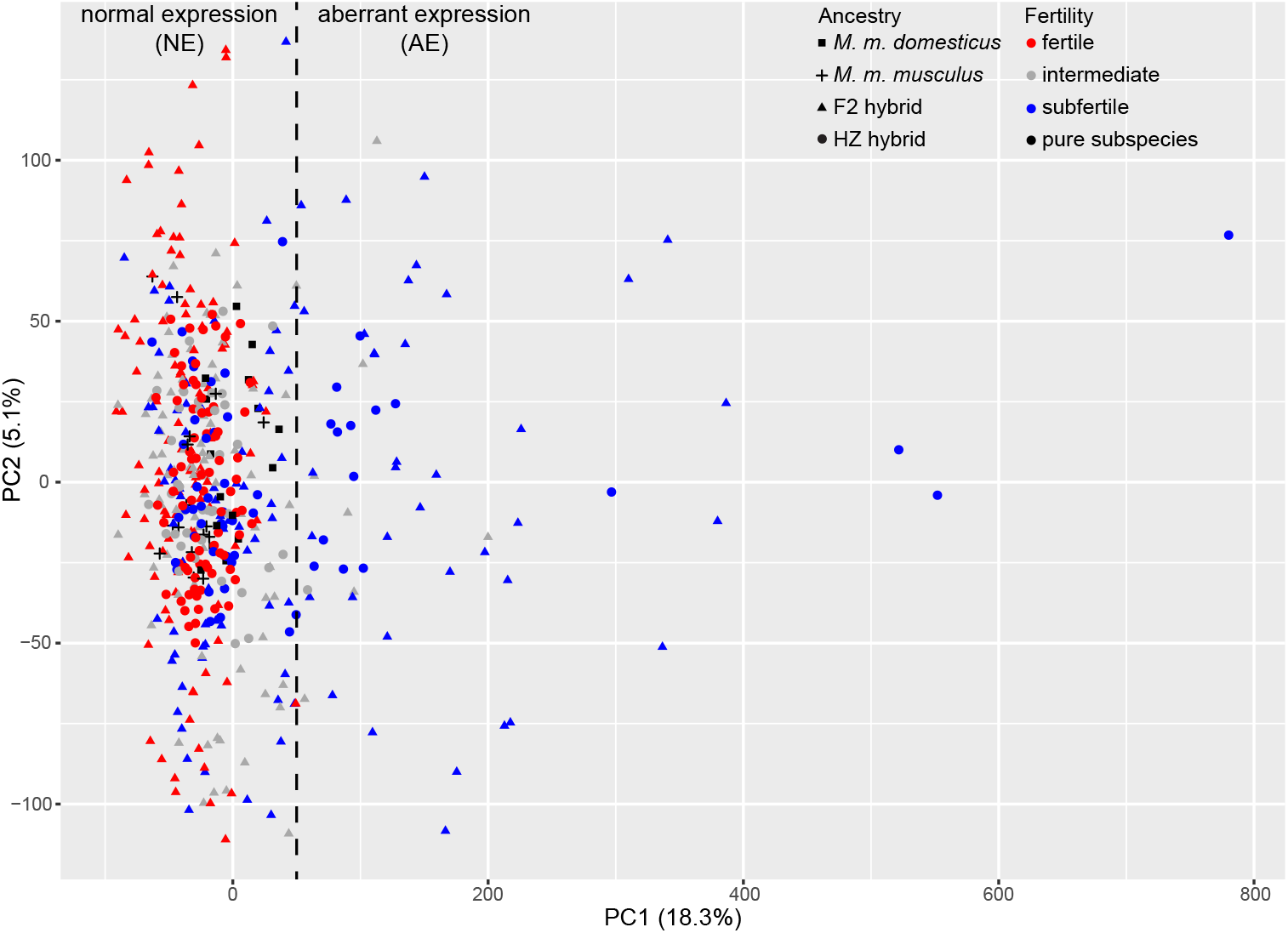
Principal components analysis (PCA) of genome-wide expression in testis of pure *Mus musculus domesticus*, M. *m. musculus* and hybrids, PC1 vs. PC2. Point shape indicates subspecies or hybrid mapping population for each individual and point colour indicates fertility class (see Methods). Subfertile hybrids with PC1 scores outside the range observed in pure subspecies males and fertile hybrids are classified as “Subfertile Aberrant Expression” (SFAE), while subfertile hybrids within the pure subspecies and fertile hybrid range were classified as “Subfertile Normal Expression” (SFNE). The dashed line indicates the cut-off between SFNE and SFAE hybrid groups.

A sizeable proportion (33.5%) of the subfertile hybrids have PC1 scores that exceed the fertile hybrid and pure subspecies range (Figure 1); we classified these individuals as “Subfertile Aberrant Expression” (SFAE), and classified subfertile hybrids that cluster with fertile hybrids and pure subspecies males along PC1 as “Subfertile Normal Expression” (SFNE). A total of 69 F_2_ and 38 HZ hybrids were categorized as SFNE, while 37 F_2_ and 17 HZ hybrids were categorized as SFAE (Figure 1). PC3, which explains 4.28% of variation, is associated with subspecies ancestry: *musculus* individuals have high scores, *domesticus* individuals have low scores, and hybrids have intermediate scores (Supplementary Figure 2).

### Fertile hybrid consensus network

To identify potentially interacting sets of genes that are coexpressed in fertile hybrids from different genomic backgrounds, we constructed a consensus fertile network using expression data from the fertile F_2_ (n=102) and HZ (n=79) hybrids and 18,411 probes representing 10,171 genes (see Methods for details). A total of 14,346 probes, representing 7,989 unique genes, were assigned to one of 15 co-expression modules (Figure 2A; Supplementary Data 1); 4,065 probes could not be assigned to a module and are shown in the grey, ‘bin’ module. Thirteen modules are significantly enriched for specific functions on the basis of gene ontology analysis, of which seven were significantly enriched for spermatogenesis or potentially related functions (Table 1, Supplementary Data 2). The module eigengene (ME), which describes the overall expression level of each module across the full dataset of fertile and subfertile hybrids, was significantly positively correlated with both sperm count and relative testis weight for three modules, and significantly positively correlated with relative testis weight alone for an additional three modules (Figure 2B). Four modules have an ME which is significantly negatively correlated with both sterility phenotypes, while the ME of one additional module is significantly negatively correlated with relative testis weight alone. Of the seven modules that are significantly enriched for spermatogenesis or potentially related functions, two (Brown and Tan) have an ME which is significantly positively correlated with at least one of the fertility phenotypes, while two (Green and Pink) have an ME which is significantly negatively correlated with at least one of the fertility phenotypes (Figure 2B). These correlations indicate that the inferred modules of coexpression are informative about fertility.

**Fig 2.**
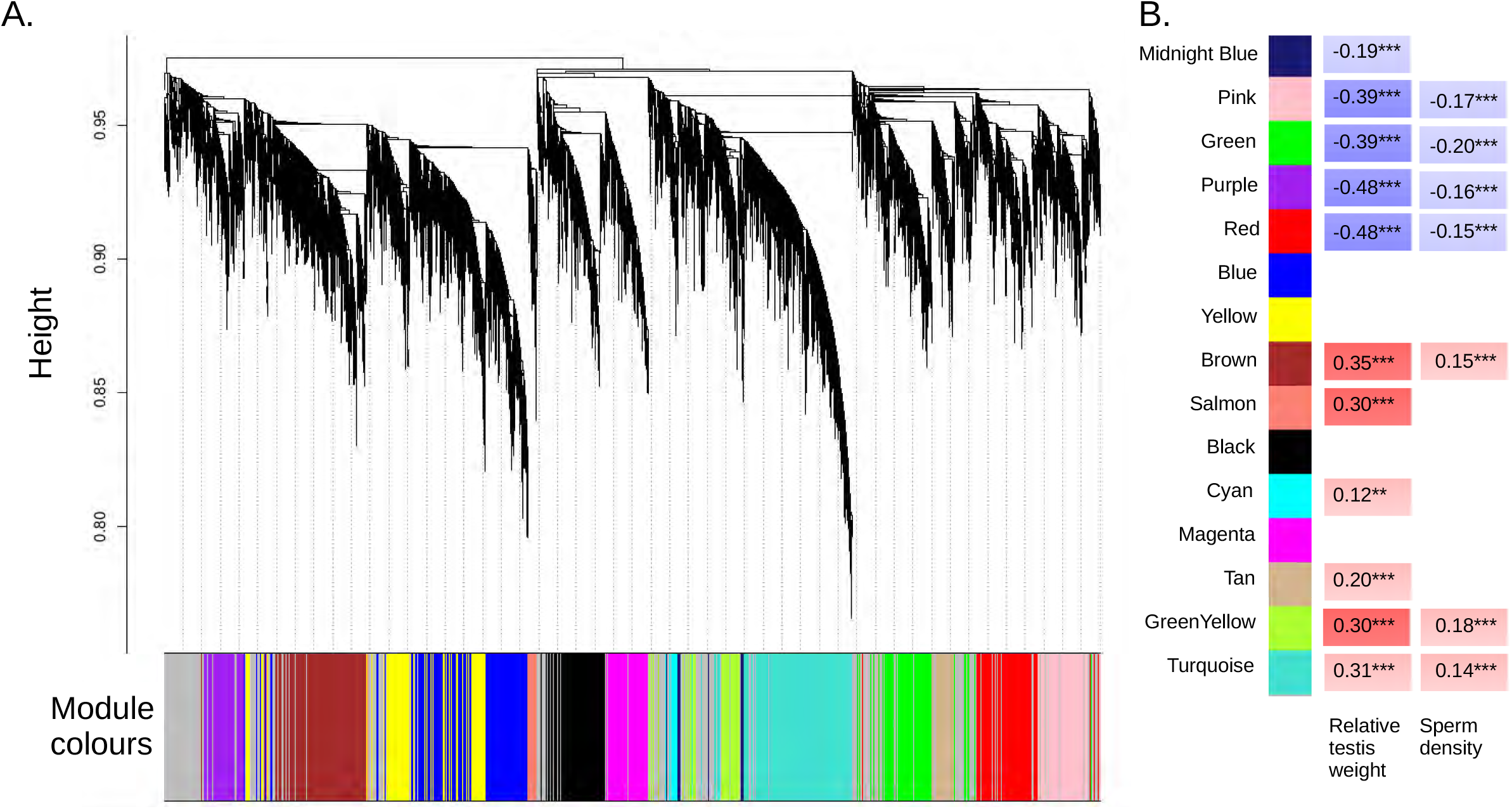
Gene co-expression modules. (A) Consensus fertile network generated using weighted gene coexpression network analysis (WGCNA; Langfelder and Horvarth, 2008) of testis expression from 102 fertile F_2_ and 79 fertile HZ hybrid males. The dendrogram shows the clustering of probes based on the topological overlap distance within fertile hybrids. Colour bar beneath the dendogram indicates coexpression modules. (B) The correlation between the Module Eigengene (ME), representing overall module expression, and sterility phenotypes. Significant positive correlations are indicated in red and significant negative correlations are indicated in blue; **p<0.01, ***p<0.001.

**Table 1.**
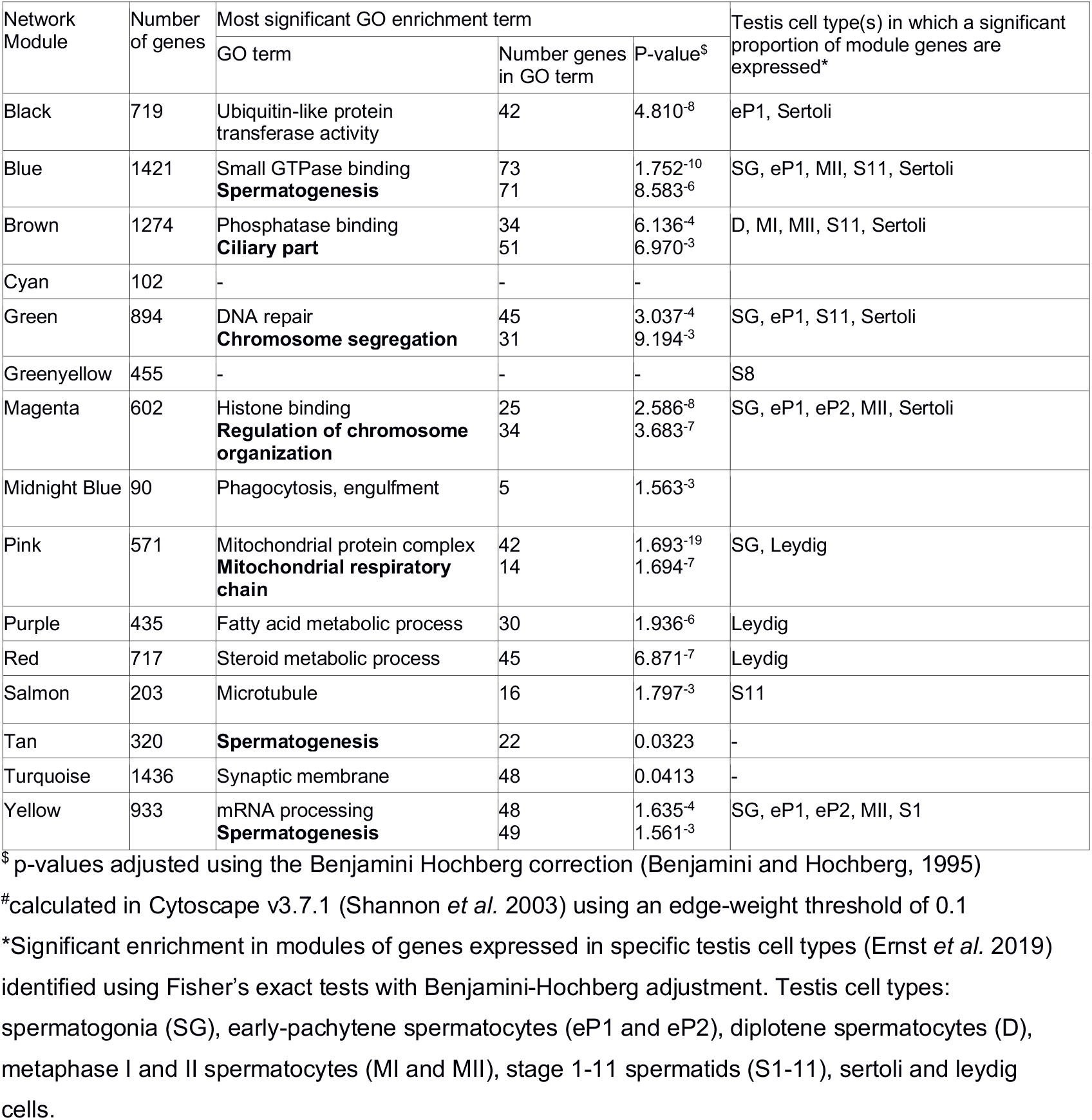
Module enrichment and connectivity within the fertile F_2_ and HZ networks. GOs potentially related to spermatogenesis are listed in bold.

### Network preservation in subfertile hybrids

To determine whether coexpression networks are disrupted in subfertile hybrids, we estimated module preservation using two approaches based on a set of metrics developed by Langfelder *et al*. (2011; see Methods). Figure 3A shows the estimated preservation of each module from the consensus fertile network in F_2_ and HZ hybrids with subfertile phenotypes and either aberrant or normal overall expression patterns. Module preservation was assessed independently for F_2_ and HZ hybrids, because the presence and prevalence of specific DMIs and associated network disruptions may vary between mapping populations. Modules showing significant evidence for preservation in subfertile hybrids are represented by circles, while modules showing a lack of significant preservation are represented by squares (Figure 3A). Figure 3B and 3C illustrate coexpression patterns within a well-preserved (Red) versus a poorly-preserved (Brown) module in the F_2_ SFAE hybrids. Pairwise coexpression in the Red module is characterized by strong positive correlations (Figure 3B). In contrast, many pairwise expression correlations in the Brown module are weakened or even reversed in direction (Figure 3C), suggesting substantial disruption in the coexpression of these gene pairs.

**Fig 3.**
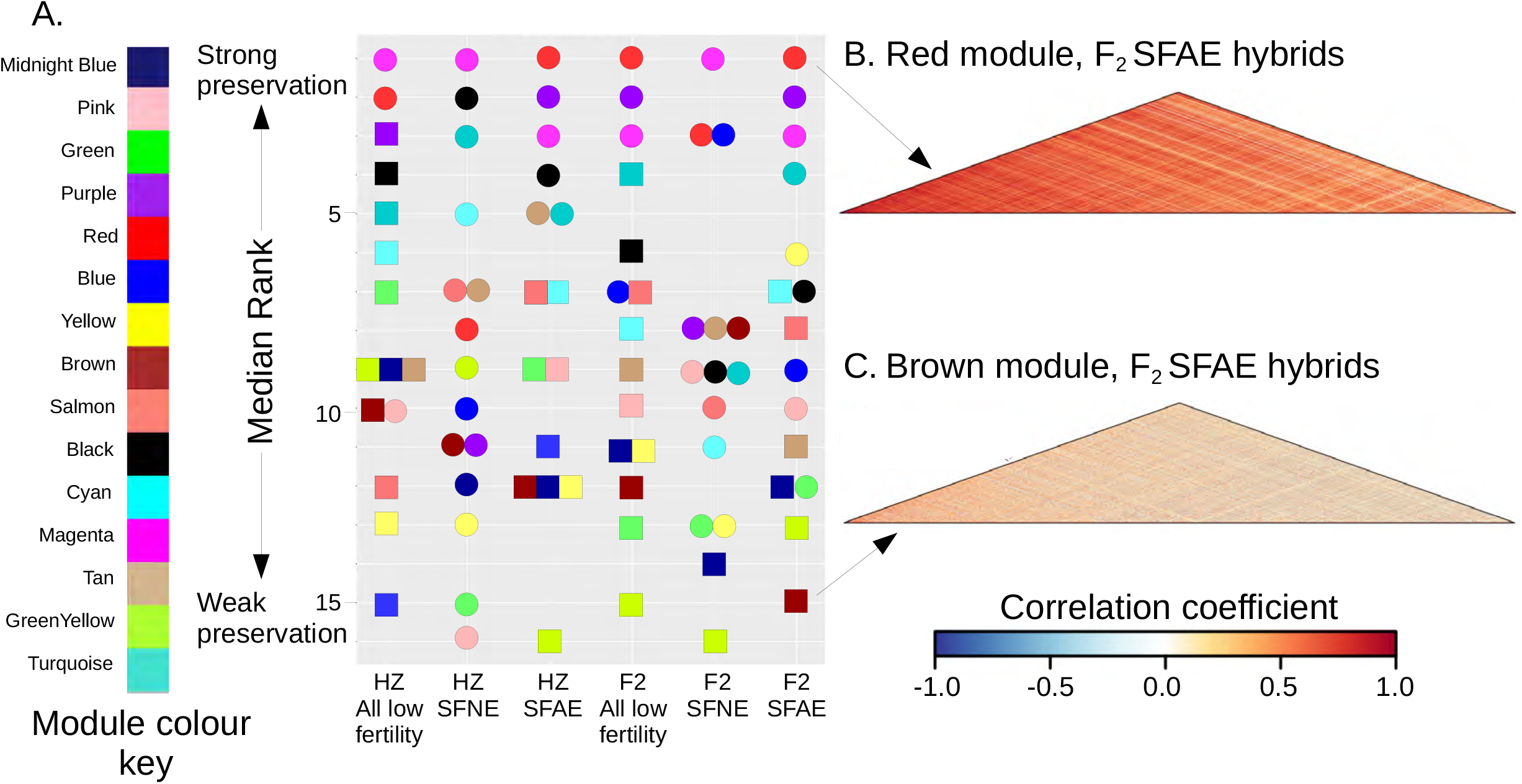
Preservation of the fertile consensus network within subfertile hybrids, classified according to mapping population derived and expression profile (SFNE or SFAE, see Figure 1). (A) Module preservation estimated using Median Rank statistics. Colour indicates module identity. Circles represent significantly preserved modules, and squares represent modules not significantly preserved. (B) and (C) Coexpression heatmaps for examples of well preserved (Red) and poorly preserved (Brown) modules. Heatmaps show pairwise correlations between expression values for all genes within modules for SFAE F_2_ hybrids.

The level of preservation of modules from the fertile network was similar in subfertile HZ and F_2_ hybrids. Three modules (Magenta, Red, Pink) showed strong evidence for preservation in subfertile HZ hybrids (Figure 3A; Supplementary Table 1); the remaining 12 modules had either a Z_summary_ < 10 and/or non-significant NetRep statistics, indicating weak or a lack of preservation (Langfelder *et al*., 2016; Ritchie et al 2016, see Methods). In F_2_ hybrids, four modules (Magenta, Red, Purple and Blue) showed strong evidence for preservation in subfertile mice.

As expected, module preservation was much higher in subfertile hybrids with normal expression based on PC1 (SFNE); all modules showed strong evidence of preservation in the HZ SFNE hybrids, and 13 of 15 modules were strongly preserved in F_2_ SFNE hybrids (Greenyellow and Midnightblue modules had non-significant NetRep scores; Supplementary Table 1). By contrast, preservation was lower in subfertile hybrids with aberrant expression; six of 15 modules were preserved in HZ SFAE hybrids, and nine modules in F_2_ SFAE hybrids.

Many modules show consistent levels of preservation across subfertile hybrid classes (Figure 3A). The Red, Magenta, Purple and Turquoise modules consistently rank among the most strongly preserved modules; these modules are significantly enriched for cofactor metabolic processes, histone binding and chromosome organization, fatty acid metabolic processes, and synaptic membrane expression, respectively (Table 1). By contrast, the Brown, Green, Yellow, Midnightblue and Greenyellow modules consistently rank among the modules with the weakest preservation; these modules are significantly enriched for cell projection assembly and cilium organization, DNA repair and chromosome segregation, mRNA processing and spermatogenesis, and protein homooligomerization (the Greenyellow module has no significant enrichments). Notably, while the Magenta and Green modules are significantly enriched for similar processes (chromosome organization and segregation, respectively), the Magenta module appears to be one of the best-preserved modules, while the Green module is among the least preserved across hybrid groups (Figure 3A).

While consistencies across subfertile groups are apparent, notable differences were also detected between the HZ and F_2_ hybrids. For example, the Blue module, enriched for spermatogenesis, is significantly preserved in the F_2_ SFAE hybrids yet shows relatively poor preservation in the HZ SFAE hybrids (Figure 3A). The Yellow module, which is also significantly enriched for spermatogenesis, is amongst the least preserved modules in all subfertile hybrid groups except for the F_2_ SFAE, within which it is relatively well preserved.

In summary, broad similarity of module preservation statistics across subfertile classes provides further evidence that specific functions and pathways are commonly disrupted in sterile hybrids. By contrast, modules showing differences in conservation in subfertile HZ vs F_2_ hybrids suggest some network disruptions are unique to specific mapping populations.

### Modules associated with specific stages of spermatogenesis or testis cell types

To determine if coexpression modules are associated with specific stages of spermatogenesis, we tested for significant enrichment of genes expressed in different testis cell types, which was recently determined at high resolution using a combination of single-cell RNAseq and bulk RNAseq at different time points during the first stage of spermatogenesis (gene lists from Supplementary Data 2 in Ernst *et al*. 2019). The gene content of five modules (Blue, Green, Magenta, Pink and Yellow) overlaps significantly with genes expressed in spermatogonia (Table 1). Six modules (Black, Blue, Brown, Green, Magenta and Yellow) overlap with genes expressed in spermatocytes during stages of meiosis (early pachytene, diplotene, metaphase). Five modules (Blue, Brown, Greenyellow, Salmon, Yellow) overlap with genes expressed postmeiotically in at least one of the 11 stages of developing spermatids. For somatic cell types, six modules (Black, Blue, Brown, Green, Magenta, Red) overlap with genes expressed in Sertoli cells and three modules (Pink, Purple and Red) are enriched for genes expressed in Leydig cells. Thus, of the 15 modules, three overlap significantly with genes expressed during spermatogenesis only (Greenyellow, Salmon and Yellow), six modules overlap significantly with both genes expressed in spermatogenesis and with genes expressed in somatic cells (Black, Blue, Brown, Green, Magenta and Pink), two modules overlap only with somatic cells (Purple and Red), and four modules do not overlap with genes expressed in any of specific testis cell types (Cyan, Midnightblue, Tan and Turquoise). Complete lists of genes with cell type-specific expression (Ernst *et al*. 2019) are in Supplementary Data 3. Our findings indicate that several coexpression modules can be linked with one or more stages of the spermatogenesis process.

### Modules associated with trans eQTL hotspots

Turner *et al*. (2014) identified 11 *trans* expression Quantitative Trait Loci (eQTL) hotspots, that is, regions of the genome with significant effects on the expression of hundreds to thousands of Quantitative Trait Transcripts (QTTs) in F_2_ hybrids, and provided several lines of evidence linking these hotspots to sterility phenotypes. We investigated whether *trans* eQTL hotspots affected specific parts of the gene coexpression network by testing for overlap between QTT associated with each hotspot and genes in modules from the fertile network. Genes in seven modules overlap with QTT from 1-7 hotspots (Table 3). Modules negatively correlated with relative testis weight and/or sperm count (Green, Red, Purple) overlap significantly with QTT showing high expression associated with the sterile allele, as established from expression patterns in sterile F_1_ hybrids (Turner *et al*. 2014), while modules positively correlated with fertility phenotypes (Brown, Greenyellow, Turquoise) overlap QTT with low expression associated with the sterile allele. That is, in all cases, ‘sterile’ vs. ‘fertile’ patterns were consistent between coexpression modules and QTT associated with sterile vs. fertile alleles.

Modules with higher preservation in subfertile hybrids (Red, Purple) overlap with QTT associated with multiple *trans*-eQTL hotspots (Table 2), highlighting potential interactions between eQTL hotspots. In contrast, modules with intermediate (Black, Turquoise) or weak (Brown, Green and Greenyellow) preservation in subfertile hybrids each overlap significantly with QTT associated with a single *trans*-eQTL hotspot. In summary, five modules with intermediate or weak preservation overlap significantly with QTT associated with a specific *trans*-eQTL hotspot, supporting an influence of the underlying sterility alleles on specific parts of the fertile gene network.

**Table 2.** Significant overlap between genes within coexpression modules and quantitative trait transcripts (QTT) associated with *trans* eQTL hotspots (Turner *et al*. 2014).

| Coexpression module | *trans* eQTL Hotspot, direction* | Sterile expression pattern<sup>#</sup> | Number of genes in module | Number of QTT | Overlap between gene sets | Fisher's exact p-value | Corrected p-value<sup>\$</sup> |
| --- | --- | --- | --- | --- | --- | --- | --- |
| Black | 18-38 cM Chr 15, Mhigh | low | 719 | 511 | 62 | 1.1e-05 | 5.1e-05 |
| Brown | 38-44 cM Chr 3, Dhigh | low | 1271 | 102 | 30 | 3.2e-06 | 4.8e-05 |
| Green | 18-38 cM Chr 15, Dhigh | high | 894 | 867 | 102 | 7.2e-04 | 0.0036 |
| Greenyellow | 18-38 cM Chr 15, Mhigh | low | 455 | 511 | 47 | 1.2e-06 | 8.4e-06 |
| Red | 26-38 cM Chr 2, Dhigh | high | 717 | 347 | 85 | 7.1e-26 | 1.1e-24 |
|  | 54-62 cM Chr 11, Dhigh | high | 717 | 763 | 166 | 5.1e-44 | 7.1e-43 |
|  | 18-38 cM Chr 15, Dhigh | high | 717 | 246 | 59 | 1.1e-17 | 8.5e-17 |
|  | 46-50 cM Chr 15, Dhigh | high | 717 | 867 | 293 | 3.5e-140 | 5.2e-139 |
|  | 0-16 cM Chr 17, Mhigh | high | 717 | 100 | 31 | 4.7e-13 | 4.7e-13 |
|  | 0-42 cM X Chr, Mhigh | high | 717 | 1117 | 171 | 3.3e-25 | 2.5e-24 |
| Purple | 26-38 cM Chr 2, Dhigh | high | 435 | 347 | 36 | 5.1e-07 | 3.8e-06 |
|  | 4-24 cM Chr 10, Dhigh | high | 435 | 763 | 88 | 7.7e-19 | 5.4e-18 |
|  | 54-62 cM Chr 11, Dhigh | high | 435 | 246 | 58 | 5.2e-28 | 8.8e-27 |
|  | 18-38 cM Chr 15, Dhigh | high | 435 | 867 | 130 | 1.0e-40 | 7.5e-40 |
|  | 46-50 cM Chr 15, Dhigh | high | 435 | 100 | 14 | 7.4e-05 | 5.55e-04 |
|  | 0-16 cM Chr 17, Mhigh | high | 435 | 1117 | 155 | 3.3e-45 | 5.0e-44 |
|  | 0-42 cM X Chr, Mhigh | high | 435 | 3329 | 238 | 6.2e-23 | 4.65e-22 |
| Turquoise | 18-38 cM Chr 15, Mhigh | low | 1434 | 511 | 115 | 4.8e-08 | 6.72e-07 |

\*Chromosome and cM position for *trans* eQTL hotspots, as reported in (Turner *et al.* 2014), and subset of QTTs associated with either the *domesticus* (Dhigh) or *musculus* (Mhigh) eQTL allele.
<sup>#</sup>QTTs showed higher or lower expression associated with the 'sterile' eQTL allele (classified on the basis of multiple criteria, see Turner *et al.* 2014 Table 2).
<sup>\$</sup>Benjamini-Hochberg correction for multiple tests.

### Differentially correlated genes

Our next aim was to identify specific genes causing observed patterns of network disruption in subfertile hybrids. We identified genes with significant changes in coexpression pattern in subfertile vs fertile hybrids using differential correlation analysis, performed independently for each module using the DGCA R package (McKenzie *et al*. 2016). A total of 2800 genes showed a significant loss or reversal of coexpression pattern within at least one of the subfertile groups (Supplementary Data 1). The percentage of differentially correlated genes varied across subfertile classes: 14.83% F_2_ SFNE, 11.08% F_2_ SFAE, 3.06% HZ SFNE, 2.88% HZ SFAE; note, these values are not necessarily consistent with degree of sterility, because sample size is smaller for HZ vs. F_2_ hybrids, and for SFAE vs. SFNE hybrids. As expected, a higher proportion of genes were differentially correlated in weakly preserved (28.89-46.59%) vs. strongly preserved (1.12-3.65%) modules (Supplementary Table 2).

### Module hub genes

We next identified hub genes in the fertile network. Hub genes are highly connected within modules, and thus more likely to be associated with network disruptions. We identified 281 hub genes across the 15 modules on the basis of degree (number of connections) and module membership (correlation between the expression of a gene and the module eigengene) (Horvath and Dong, 2008). Of the 281 hub genes, 95 genes within 10 modules show a significant loss or reversal of coexpression pattern in at least one of the subfertile hybrid groups relative to the fertile hybrids (Table 3). Of those 94 differentially correlated hub genes, 29 and 57 show a significant loss of coexpression pattern in the SFAE and SFNE F_2_, respectively, while 11 and 10 show a significant loss of coexpression pattern the SFAE and SFNE HZ, respectively (Supplementary Data 4). Hence, although the WGCNA and NetRep analyses show evidence for stronger module preservation in SFNE relative to SFAE hybrids, differential correlation analysis shows evidence for the disruption of module hub gene interactions in both the SFAE and SFNE hybrid groups.

**Table 3.**
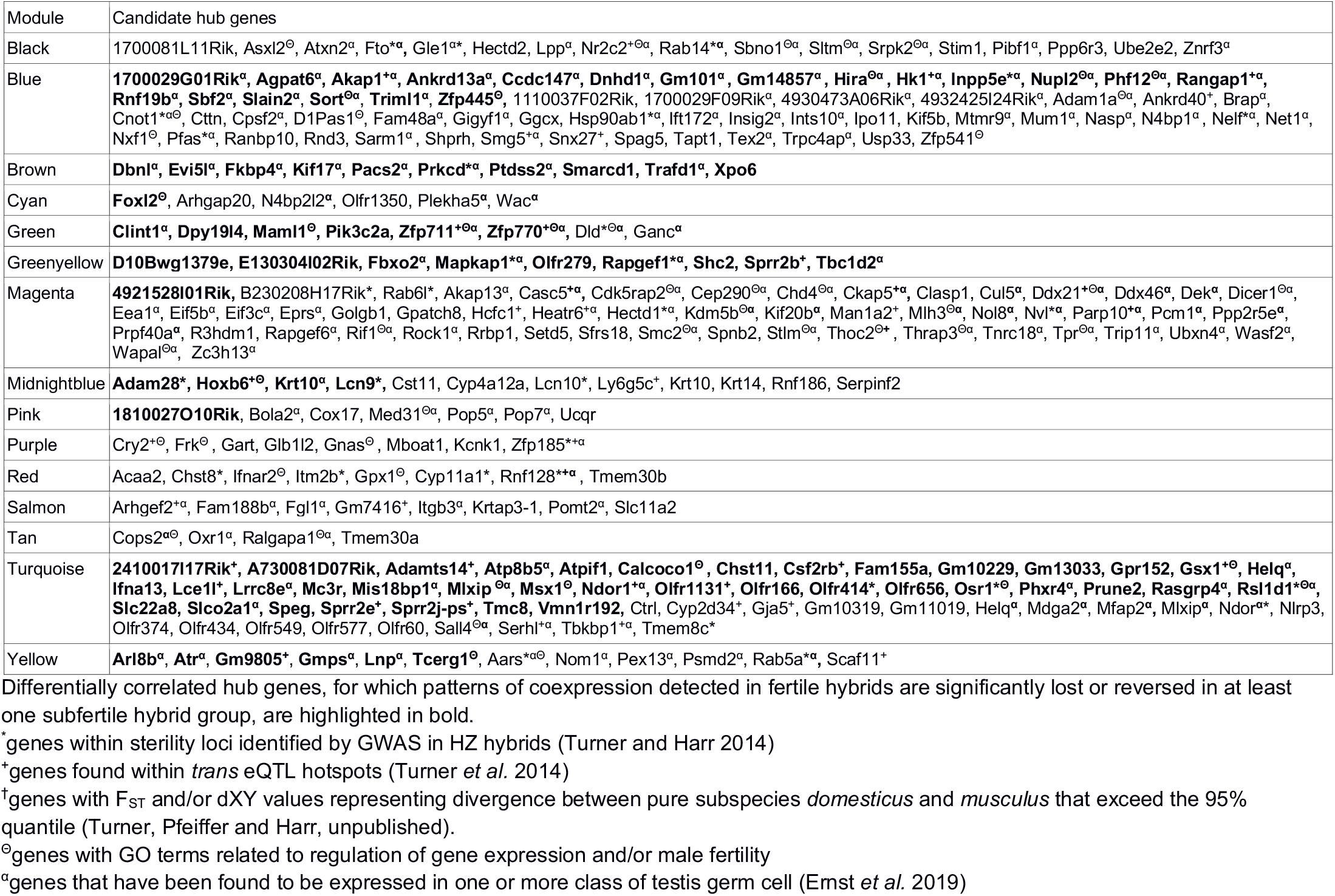
Candidate module hub genes.

Table 3 lists the module hub genes, indicating the following as potential candidates for DMIs: genes with different coexpression patterns in subfertile relative to fertile hybrids (n=95); genes that have been found to be expressed within spermatogonia, spermatocytes or spermatids (Ernst *et al*. 2019) (n=152); genes with GOs including those related to the regulation of gene expression and/or male fertility (n=52); genes that fall within previously identified *trans* eQTL hotspots (Turner *et al*. 2014) (n=40); genes within sterility loci identified by GWAS in HZ mice (Turner and Harr 2014) (n=31).

All hub genes in the Brown module are differentially correlated in at least one subfertile hybrid group relative to fertile hybrids. Figure 4 illustrates the widespread loss of connectivity within the SFAE relative to the fertile F_2_ network. The loss of interactions for four of the module hub genes, *Prkcd, Dbnl, Ptdss2* and *Fkbp4*, are shown in Figure 4A, and the overall pattern of reduced connectivity in the SFAE network is shown in Figure 4B. Several genes that interact with module hub genes in the fertile hybrid network, including *Spata18, Ttll5, Gli2, Sept2* and *Adcy3*, have Cell Component (CC) GOs that include cilium and/or sperm flagellum (Figure 4A), which is of note since the Brown module is significantly enriched for genes involved in cilium organization (Table 1).

**Fig 4.**
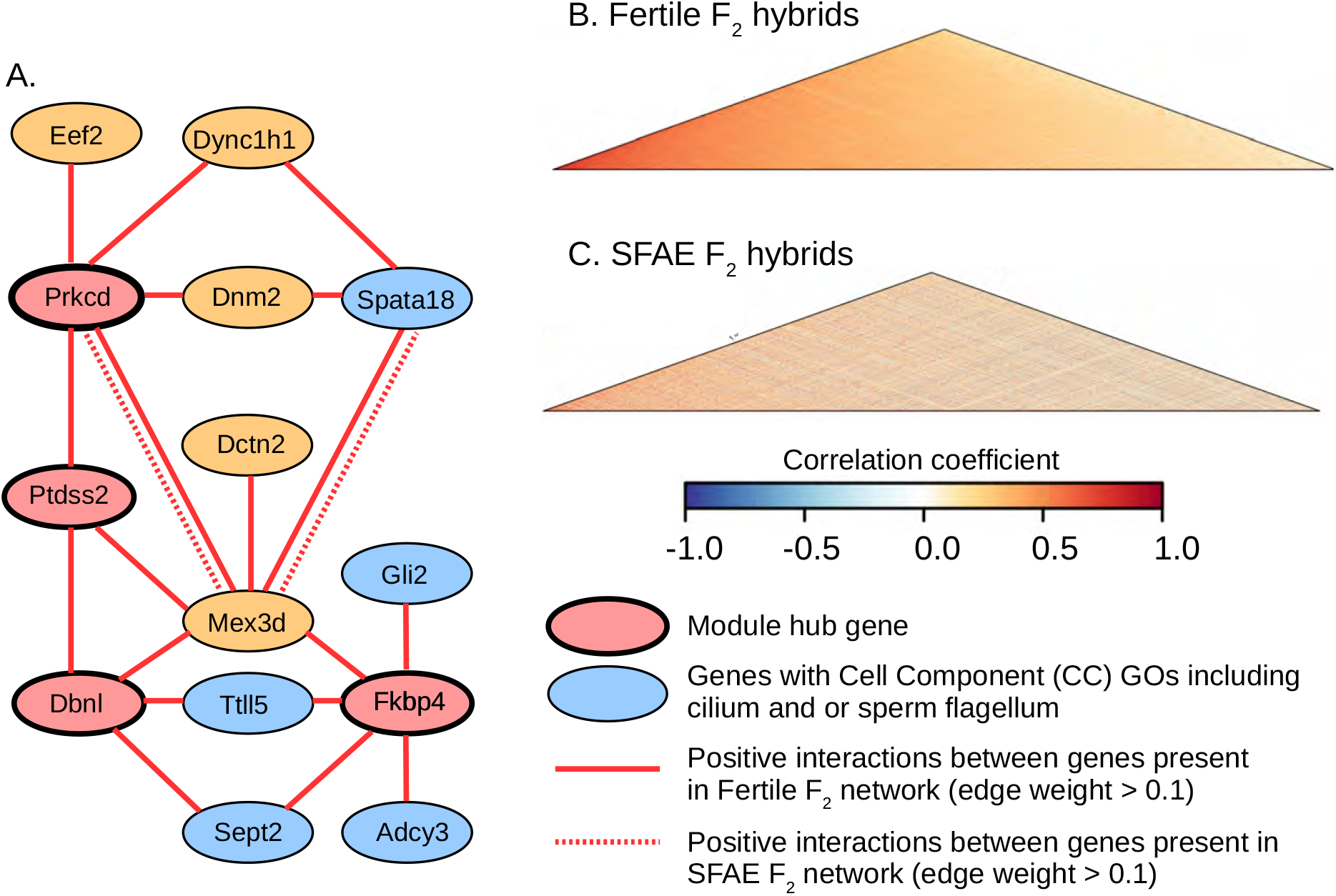
Disrupted interactions of Brown module genes in subfertile F_2_ hybrids. (A) Interactions between Brown module hub genes (red nodes) and genes with GOs including cilium/and or sperm flagellum (blue nodes) in fertile and SFAE F_2_ hybrids. Orange nodes indicate intermediate genes with functions potentially related to male fertility. Gene interactions with an edge-weight exceeding 0.1, as estimated using topological overlap matrices, are indicated using continuous and dashed lines for the fertile and SFAE hybrids, respectively. (B) and (C) Coexpression heatmaps showing pairwise correlations between expression values of Brown module genes in fertile and SFAE F_2_ hybrids, respectively.

We found some hub genes in significantly preserved modules that nevertheless show disrupted coexpression patterns in subfertile hybrids. For example, the Blue module is significantly preserved in the SFAE F_2_ hybrid group (Figure 3), however several hub genes show significant loss of interactions in SFAE F_2_ hybrids, including *Hk1, Akap1, Agpat6* and *Slain2* (Table 3; Supplementary Figure 3A; Supplementary Data 4). The Blue module coexpression heatmaps for fertile and SFAE F_2_ hybrids (Supplementary Figure 3B-3C) show weakening of positive correlations in these subfertile hybrids. Meanwhile, the Midnightblue module was preserved in SFNE HZ hybrids, despite showing a lack of preservation in all other subfertile groups (Figure 3). However, two hub genes in this module have changes in interactions in both SFNE and SFAE HZ hybrids (Figure 5A). Moreover, an intermediate level of reduced connectivity overall in SFNE HZ hybrids in the Midnightblue module is apparent from visual comparison of the coexpression heatmap of the module compared to fertile hybrids and the more severely disrupted SFAE hybrids (Figure 5B-D).

**Fig 5.**
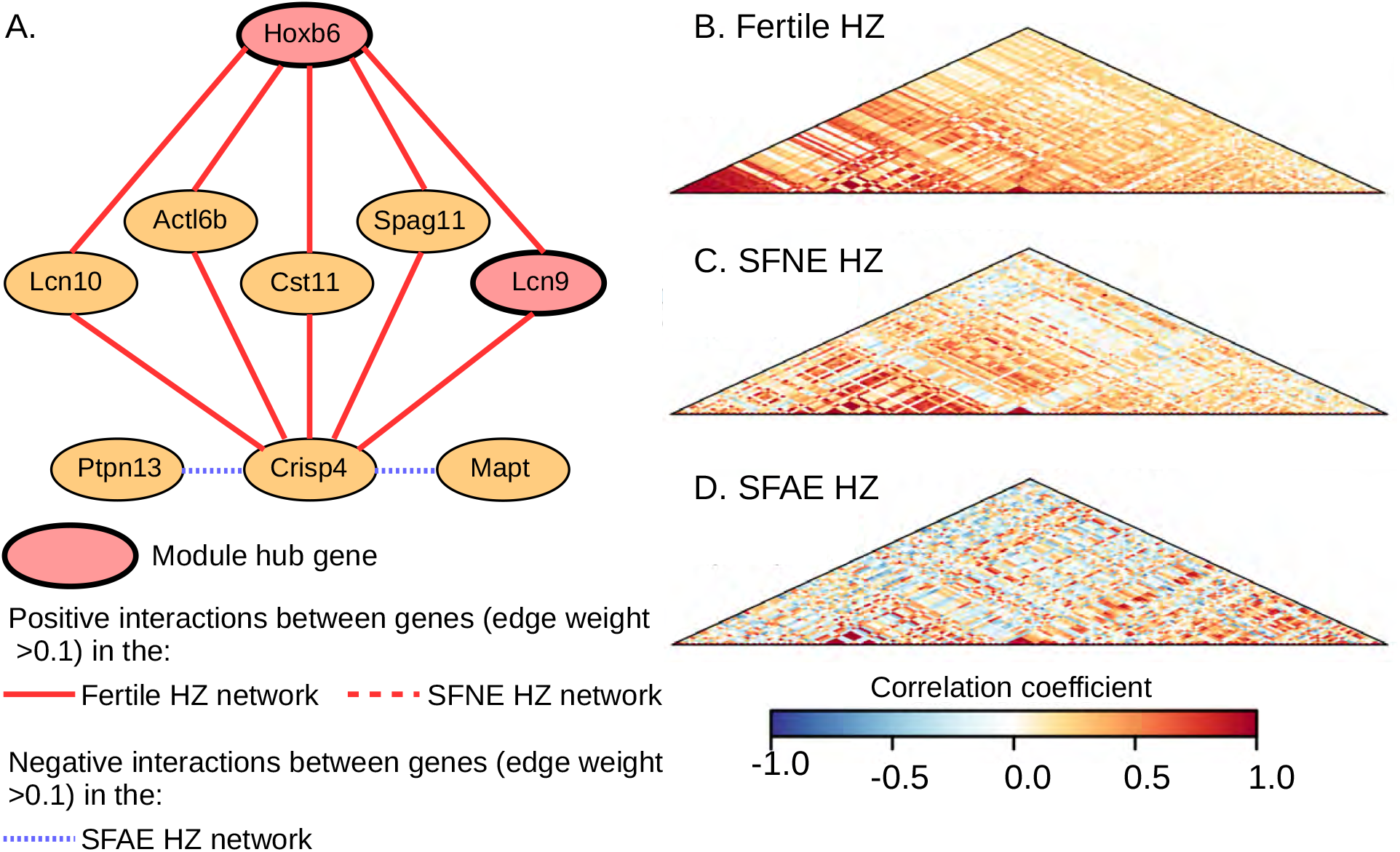
Disrupted interactions of Midnightblue module genes in subfertile HZ hybrids. (A) Interactions between Midnightblue module hub genes (red nodes) and genes with functions potentially related to sperm maturation (orange nodes) in HZ hybrids. Gene interactions with an edge-weight exceeding 0.1, as estimated using topological overlap matrices, are indicated using continuous, dashed and fine-dashed lines for the fertile, SFNE and SFAE hybrids, respectively. Positive interactions are shown in red and negative interactions are shown in blue. (B), (C) and (D) Coexpression heatmaps showing pairwise correlations between expression values of Midnightblue module genes in fertile, SFNE and SFAE HZ hybrids, respectively. Heatmaps reveal moderate weakening of gene interactions in the SFNE hybrids and weakening or reversal of interactions in the SFAE HZ hybrids, indicating more severe network disruption in the SFAE hybrid group.

## Discussion

While the Dobzhansky-Muller model provides a well-accepted mechanism for the development of reproductive isolation between diverging lineages, the specific epistatic interactions underlying DMIs remain mostly uncharacterized. A network approach is ideal for identifying complex DMIs, since pairwise interactions between genes are likely to be non-independent. Here, using hybrids of two house-mouse subspecies between which reproductive barriers are incomplete (Britton-Davidian *et al*. 2005; Albrechtová *et al*. 2012; Turner *et al*. 2012), we take a network-based approach to characterize gene interactions in testis of fertile vs. subfertile males from two hybrid mapping populations. Combining testis expression data from an F_2_ cross between wild-derived inbred strains (Turner *et al*. 2014) and offspring of mice wild-caught from a hybrid zone (Turner and Harr 2014) enabled us to generate a consensus fertile network and identify modules of genes that are commonly coexpressed. We identified disruptions in this network in subfertile hybrids at the module and gene level, some of which are associated with specific functions, stages of spermatogenesis, or testis cell types. Integrating our results with previous mapping of subfertility phenotypes and gene expression traits in the same mice (White *et al*. 2011; Turner *et al*. 2014; Turner and Harr 2014) reveals candidate pathways and genes for subfertility.

### Network disruptions in subfertile hybrids

Several modules show consistent patterns of poor preservation in subfertile hybrids within both the F_2_ and HZ mapping populations, including modules enriched for cilium organization and the sperm flagellum (Brown), spermatogenesis and the regulation of cell cycle (Blue and Yellow), DNA repair and chromosome segregation (Green). The disrupted modules are enriched for genes expressed in spermatogonia (Blue, Green and Yellow), spermatocytes (Blue, Brown, Green and Yellow) and spermatids (Blue, Brown, Greenyellow and Yellow). Hence, our findings suggest that multiple stages of spermatogenesis are impacted by DMIs in both natural and laboratory-bred hybrid mice. These findings are somewhat supported by previous histological analyses of testis defects in the F_2_ hybrid mapping population, which revealed a range of phenotypic defects linked to reduced fertility (Schwahn *et al*. 2019). While the majority of these defects could be explained by a failure to complete meiosis I, so potentially implicating genes expressed in primary spermatocytes, possible downstream errors in meiosis II and postmeiotic errors in spermiogenesis were also implicated (Schwahn *et al*. 2018).

Although most modules enriched for functions potentially related to spermatogenesis show a lack of preservation in at least one of the subfertile hybrid groups, the Magenta module is an exception. This module is relatively well-preserved in all subfertile hybrid groups, despite being enriched for similar processes to the poorly preserved Green module (chromosome organization) and despite being enriched with genes expressed in spermatogonia and spermatocytes. This pattern suggests that while several different stages of spermatogenesis are potentially impacted by DMIs, some pathways and processes remain intact.

Patterns of network disruption are broadly similar in the F_2_ and HZ mapping populations, but differences are also evident. The Blue and Yellow modules, for example, showed higher levels of preservation in the F_2_ relative to HZ subfertile hybrids, and several module hub genes showed significant changes in coexpression pattern in only one of the hybrid populations. As noted above, power to detect disruptions varies among subfertile classes, due to sample size, but we expect there are also true biological differences, because incompatibility loci are segregating within *musculus* and *domesticus* (Good et al. 2008b; Larson *et al*. 2018). Moreover, the HZ mapping population is more genetically and phenotypically diverse (Turner et al. 2012), hence the specific genes driving network disruptions may vary across hybrid populations. It is possible that severe DMIs might have been purged by selection in hybrid zone, and are consequently detectable in F2 but not HZ hybrids. However, there is ongoing gene flow from pure subspecies populations, and most sterility phenotypes appear to have modest fitness effects. Hence, selection against incompatibilities in the hybrid zone seems unlikely to be a general pattern explaining polymorphism. However, the lack of a prominent role of *Prdm9* in sterility of Bavarian hybrid-zone mice is consistent with this hypothesis.

### Variation in patterns of network disruption

We split hybrid mice with subfertile phenotypes into two broad categories: mice with similar overall patterns of gene expression to those seen in fertile hybrids (SFNE) and those with relatively aberrant patterns of gene expression (SFAE). Unsurprisingly, network disruption appears more severe in the SFAE hybrid groups from both the F_2_ and HZ populations, with fewer modules showing evidence of significant preservation. However, there is evidence for more subtle network disruption in the SFNE hybrids. Two modules are not preserved in the SFNE F_2_ hybrid group (Midnightblue and Greenyellow, Figure 3). In SFNE HZ hybrids, all modules are preserved but intermediate levels of disruption are apparent upon detailed examination of module-specific coexpression patterns within the Midnightblue module (Figure 5).

We identified more fine-scale disruptions in networks by performing differential correlation analysis, which detects genes showing a significant change in coexpression pattern between groups. Genes showing a significant loss or reversal of coexpression patterns were detected in both SFNE and SFAE hybrid groups relative to fertile hybrids, in both the F_2_ and HZ mapping populations, supporting an influence of subtle network disruptions in all subfertile hybrid groups. The 2800 genes with different interactions in at least one subfertile group include 94 module hub genes, some of which are in relatively well-preserved modules, suggesting hybrid incompatibilities can cause minor or major perturbations in gene interactions. Hence, our findings support varying levels of network disruption within and among subfertile hybrid groups, as expected because sterility loci are segregating within the mapping populations and putative DMIs previously identified show a range of complexity and effect size (White *et al*. 2011; Turner *et al*. 2014; Turner and Harr 2014).

### Overlap with previously identified sterility regions

Several modules, including the Black, Brown, Green, Greenyellow, Red, Purple and Turquoise modules, have gene content that overlaps significantly with QTT that are associated with a specific *trans*-eQTL hotspot. The majority of these modules show weak (Brown, Green, Greenyellow) or intermediate (Black, Turquoise) preservation in subfertile hybrids, potentially supporting a role for network disruptions involving specific *trans*-eQTL. The Purple and Red modules, however, are well preserved across subfertile hybrid groups and overlap significantly with QTT associated with multiple *trans*-eQTL hotspots. Unlike the more weakly preserved modules, very few of the Purple and Red module genes are expressed during spermatogenesis (Ernst *et al*. 2019), rather both modules are enriched for genes expressed in Leydig cells. The overall expression of both modules is negatively correlated with both fertility phenotypes, suggesting expression tends to be higher in subfertile relative to fertile hybrids. These observations are consistent with previous reports that genes expressed in somatic cells in testis (i.e. Leydig, Sertoli) have relatively high expression in subfertile hybrids (Turner *et al*. 2014), likely reflecting reduction/absence of germ cells.

### Candidate DMI genes

Module hub genes, the most well-connected genes in the fertile network, are good candidates for large-effect DMIs, since disrupted epistatic interactions involving these genes may have knock-on effects on module-wide gene expression. We compared these module hub genes to previously identified candidate DMI loci, which fall within *trans*-eQTL hotspots (Turner *et al*. 2014) and/or GWAS sterility regions (Turner and Harr, 2014) and have been prioritized as candidates based on their Gene Ontology (GO) categories. Module hub genes previously prioritized as candidate DMI loci include *Nr2c2, Zfp711, Zfp770, Hoxb6, Cry2* and *Gsx1*, all of which lie within *trans*-eQTL hotspots, and *Cyp11a1*, which is located within a GWAS sterility locus on chromosome 9. All of these genes have GO categories that include the regulation of transcription and/or spermatogenesis. The Black module hub gene *Nr2c2* is of particular interest. *Nr2c2* is expressed in mid- and late-stage pachytene spermatocytes and round spermatids (Ernst *et al*. 2019; Mu *et al*. 2004) and a lack of expression has been associated with disruptions to late meiotic prophase and consequent delays to spermiogenesis (Mu *et al*. 2004). The transcription factors *Zfp711* (Green module), *Zfp770* (Green module), *Hoxb6* (Midnightblue module) and *Gsx1* (Turquoise module) are also good candidate loci for large-effect DMIs, since they show a loss of coexpression patterns in subfertile relative to fertile hybrids and are hub genes within modules that show weak (Green and Midnightblue) or intermediate (Turquoise) preservation in subfertile hybrids.

*Trans*-eQTL hotspots and many GWAS sterility loci each contain numerous genes (Turner *et al*. 2014; Turner and Harr, 2014), and honing in on the specific causative genes driving DMIs has been challenging. Genes that lie within candidate regions but lack GOs relating to the regulation of transcription and/or spermatogenesis are likely to have been overlooked. Our study identified several hub genes that lie within previously identified sterility regions yet have not previously been highlighted as likely candidate DMI loci. One such gene is *Prkcd*, a hub gene within the poorly conserved Brown module that lies within a GWAS sterility region on chromosome 14 (Turner and Harr, 2014). Although this gene lacks GOs related to spermatogenesis, a knockout study has reported a role for *Prkcd* in male fertility. Specifically, the sperm of male mice lacking *Prkcd* expression have a reduced ability to penetrate the zona pellucida, potentially impairing fertilization (Ma *et al*. 2015). *Hk1*, a hub gene within the Blue module, lies within a *trans*-eQTL hotspot on chromosome 10 yet also lacks GOs related to spermatogenesis. This gene encodes the enzyme that initiates the glycolysis pathway, which is important for sperm motility (Mori *et al*. 1998), hence suggesting an important role in male fertility. Other genes that lie within *trans*-eQTL hotspots and/or GWAS sterility regions yet have been overlooked as candidate DMI loci include *Rapgef1* and *Mapkap1*, which are involved in the regulation of signal transduction and establishment of actin cytoskeleton polarity, respectively, *Akap1*, which is known to be involved in meiosis in female mice (Newhall *et al*. 2006), and *Atr*, which plays a role in preventing DNA damage and is associated with reduced testis weight, abnormal DNA replication and cell cycle in knockout male mice (Murga *et al*. 2009).

While several hub genes within the poorly preserved Midnightblue module lie within previously identified sterility regions, only *Hoxb6* has been highlighted as a candidate DMI locus. *Adam28, Lcn10* and *Lcn9* all lie within GWAS sterility regions, and both *Adam28* and *Lcn9* show a loss of positive coexpression patterns in subfertile relative to fertile hybrids. While all three genes lack GOs relating to spermatogenesis, *Adam28, Lcn9* and *Lcn10* are all known to be expressed in the epididymis, and are likely to be involved in male fertility (Oh et al. 2005; Suzuki *et al*. 2014). Although our expression data are from testis; we suspect most of these genes are expressed in both tissues as testes are relatively easy to separate from epididymis during dissection. In fertile hybrids the Midnightblue module hub genes are coexpressed with several genes thought to be involved in sperm maturation, including *Cst11* and *Spag11*, both of which have antimicrobial activity and are thought to be important for sperm maturation in other mammal species (e.g. Avellar *et al*. 2007; Hamil *et al*. 2002), and *Crisp4*, which is implicated in the acrosome reaction required for the binding of sperm to the zona pellucida (Turunen *et al*. 2012). Hence disrupted interactions in Midnightblue module hub genes may have an impact on sperm motility and functioning.

Our gene network analysis is largely independent of phenotype data, hence potentially enabling us to identify entirely novel candidate DMI loci. Candidate DMI genes that lie outside of previously identified sterility regions include *Fkbp4* and *Ptdss2*, which are hub genes in the poorly preserved Brown module and show loss of coexpression patterns in subfertile relative to fertile hybrids. Mice lacking *Ptdss2* expression have reduced testis weight and can be infertile (Bergo *et al*. 2002), while the lack of *Fkbp4* expression is associated with abnormal sperm morphology (Hong *et al*. 2007). *Helq*, a differentially correlated hub gene in the Turquoise module, also lies outside of previously identified sterility regions and is associated with subfertile phenotypes in male mice (Adelman *et al*. 2013). Finally, *D1Pas1* and *Adam1a*, both hub genes in the Blue module have GOs relating to spermatogenesis and the binding of sperm to the zona pellucida, respectively, yet have not been previously identified as candidates for DMI speciation genes.

### Conclusion

We demonstrate widespread disruptions to gene-interaction networks in association with reduced fertility in hybrid *musculus-domesticus* house mice. Disruptions are variable in magnitude among hybrid mapping populations and appear to affect multiple stages of spermatogenesis, including chromosome segregation and cell cycle, assembly of the sperm flagellum, and sperm maturation. We identify specific candidate DMI genes, several of which fall within previously identified sterility loci and have been previously associated with reduced fertility phenotypes in male mice.

## Methods

### Microarray and phenotype data

We used testis gene expression data from two previous studies, the first included F_2_ hybrid males from a cross between wild-derived inbred strains of *M. m. domesticus* (WSB/EiJ) and *M. m. musculus* (PWD/PhJ) (White *et al* 2011; Turner *et al*. 2014), and the second included first-generation offspring of mice wild-caught in the hybrid zone (Turner and Harr 2014). Gene expression in the testis of hybrid males sacrificed at a similar developmental stage (70 +/- 5 days and 9-12 weeks for F_2_ and HZ mice, respectively) was measured using Whole Mouse Genome Microarrays (Agilent), as described in Turner *et al*. (2014) and Turner and Harr (2014).

To investigate changes in gene expression networks associated with sterility, we first classified individuals as ‘fertile’ vs. ‘subfertile.’ As fertility (*i.e*., ability to father offspring) was not directly measured in F_2_ or HZ individuals, we used two phenotypes to categorize males as fertile or subfertile: relative testis weight (testis weight/body weight) and sperm count (Turner *et al*. 2012; White *et al*. 2011). These phenotypes were measured comparably in mice from both mapping populations and have been associated with reduced fertility in multiple studies of *musculus-domesticus* hybrids (Britton-Davidian *et al*. 2005; reviewed in Good *et al*. (2008), (2010)); we will henceforth refer to these traits as “sterility phenotypes”. A total of 102 F_2_ and 79 HZ hybrid males have fertility phenotypes that fall within one standard deviation of the pure subspecies mean and were categorized as “fertile”. “Subfertile” hybrids, for which one or both of the sterility phenotypes fall outside of the pure subspecies range include 107 F_2_ and 55 HZ males. For a total of 92 F_2_ and 41 HZ hybrids, both sterility phenotypes fall within the pure subspecies range yet at least one of the phenotypes is more than one standard deviation from the pure subspecies mean. These hybrids were categorized as “intermediate phenotype”. Finally, data was available for 32 pure subspecies males: 16 *domesticus* and 16 *musculus*. Eight individuals each of pure *domesticus* and pure *musculus* were offspring of mice wild-caught at the edges of the hybrid zone (Turner and Harr 2014), and the remaining pure subspecies males were from wild-derived inbred strains WSB/EiJ (*domesticus*) and PWD/PhJ (*musculus*) inbred strains, whose expression was reported in Turner *et al*. (2014). In total, microarray and sterility phenotype data were available for 467 hybrid and 32 pure subspecies males.

### Microarray data processing

The Whole Mouse Genome Microarray (Agilent) contains 43,379 probes including 22,210 transcripts from 21,326 genes. We started from raw array data from each study rather than processed expression values, to ensure data sets were comparable in network analyses. Preprocessing of raw expression data was performed in the R package limma (Smyth, 2005). Background correction was performed by specifying the “half” setting, which resets intensities that fall below 0.5 following background subtraction to 0.5, and by adding an offset of 50. To identify probes with consistently low expression, the 98^th^ percentile of the expression of negative control probes was calculated and only probes that were at least 10% brighter than this background expression level were retained, reducing the dataset to a total of 36,896 probes. The Quantile method was used to normalise expression between arrays.

Since the expression dataset includes data generated within different laboratories and over different time periods, non-biological systematic bias or “batch effects” must be considered. We adjusted for known batch effects using the empirical Bayes framework implemented via the ComBat function (Johnson *et al*. 2007). We also tried detecting and adjusting for hidden batch effects using the SVA R package (Leek and Storey, 2007) and obtained similar results, with several of the detected surrogate variables clustering the data by the known batches. Adjusting for the batch effect may result in losing potential heterogeneity in gene expression between the F_2_ and HZ mapping populations. Nevertheless, this effect should be equal for both fertile and subfertile phenotypes, and should therefore have minimal impacts on observed patterns of network disruption in subfertile hybrids. Variation in gene expression across individuals was summarised using Principal Components Analysis (PCA), as implemented using the prcomp function in R (R Core Team, 2018), which uses a singular value decomposition of the centred and scaled data matrix, and extreme outliers were identified visually and removed prior to downstream analyses. Batch-corrected and normalised expression data has been submitted to the Gene Expression Omnibus (GEO).

### Weighted Gene Coexpression Network Analysis

We used The WGCNA R package (Langfelder and Horvarth, 2008) to identify groups of genes, “modules”, showing similar patterns of expression within and across the F_2_ and HZ fertile hybrid groups. Because network analysis is computationally intensive, we further filtered the dataset prior to network construction. Specifically, the connectivity of each probe was estimated using the softConnectivity function, which is available within the WGCNA R package (Langfelder and Horvarth, 2008) and calculates the sum of the adjacency of each probe to all other probes within the dataset. The median connectivity was calculated and probes with above-average connectivity within fertile hybrids were retained, resulting in a final dataset of 18,411 probes representing 10,171 genes. We used the blockwiseConsensusModules function to perform signed network construction and identify consensus modules across the fertile F_2_ and HZ datasets, assigning each probe to a single module. Briefly, this process involved calculating a pairwise coexpression matrix for each of the F_2_ and HZ fertile groups, in which coexpression is estimated using Pearson correlation values. Raising the coexpression matrix to a defined soft-threshold power introduces scale-free topology, in which a small proportion of nodes (hub genes) have a large number of connections within the network. Such scale-free topology is thought to be a fundamental property of most biological networks (Barabasi and Albert 1999). Network topology analysis was performed for a range of soft-threshold values, and an optimal soft-threshold value of five was chosen as the lowest value at which median connectivity reached a low plateau. The coexpression matrix was raised to this soft-threshold power to create an adjacency matrix, which was then converted to a Topological Overlap Matrix (TOM). The TOM describes the network interconnectivity or coexpression between each pair of genes in relation to all others in the network. A consensus TOM was then used to cluster genes using average linkage hierarchical clustering. A dynamic tree cutting algorithm was used to cut the clustering tree, so defining consensus modules of similarly expressed genes. The deepSplit and minimum module size parameters were set to 0 and 50 respectively, and the module eigengene distance threshold was set to 0.2 to merge similar modules.

Module eigengenes are defined as the first principal component describing the expression of a given module. To determine whether specific modules are more or less expressed in fertile vs. subfertile individuals, Pearson correlations were computed between the eigengene of each module and each of the two sterility phenotypic trait values (relative testis weight, sperm count).

### GO enrichment analysis

We tested for functional enrichment within each module on the basis of gene ontology (GO) over-representation analysis with Benjamini-Hochberg p-value adjustment (Boyle *et al*. 2004), performed using the enrichGO function from the clusterProfiler R package (Yu *et al*. 2012). The gene universe consisted of 21,200 genes, established using all Entrez identification numbers (henceforth “genes”) associated with the G4122F microarray (Agilent Whole Mouse Genome Microarray) probes via the Gene Expression Omnibus entry for the platform (Edgar et al., 2002; Barrett et al. 2013). GO terms associated with more than 10 and fewer than 500 genes in the gene universe were available for assignment.

### Network preservation between fertile and low fertility hybrid groups

To identify gene interactions which are present in fertile hybrids and disrupted in hybrids with low fertility, we tested for preservation of modules from the fertile network in subfertile hybrids. Levels of genetic variability are likely to vary between the HZ and F_2_ mapping populations, since the F_2_ hybrids were created through crosses of inbred *domesticus* and *musculus* strains, whereas the HZ population was created through crosses of mice caught wild in the hybrid zone. Since incompatibility loci are likely to be segregating in natural *domesticus* and *musculus* populations, the presence or absence of specific sterility loci is likely to differ between the mapping populations. We therefore tested for module preservation in subfertile hybrids independently within the F_2_ and HZ populations. Because the PCA revealed a strong association between the low fertility phenotype and variation along PC1 (see Results), we further split the subfertile individuals into those clustering together with the fertile individuals along PC1 and those with a PC1 score that falls outside of the fertile range (−91.25 – 49.33; see Figure 1). We refer to these groupings as “SubFertile Normal Expression” (SFNE) and “SubFertile Aberrant Expression” (SFAE). To explore whether the lack of preservation for several modules in the subfertile phenotypes was exclusive to the SFAE group, we tested for module preservation between the fertile and each of the SFNE and SFAE groups within the F_2_ and HZ populations.

We used the statistical frameworks implemented in the WGCNA and NetRep R packages to estimate module preservation (Langfelder and Horvarth, 2008; Ritchie *et al*. 2016). Both of these permutation-based approaches use the seven preservation metrics developed by Langfelder *et al*. (2011). The modulePreservation function (WGCNA package, 500 permutations) was used to generate Z_summary_ scores, which combine several preservation statistics that compare the density and pattern of connections within modules and between datasets. Z_summary_ scores of ≥ 10, 2-10, and <2 indicate strong, weak, and a lack of module preservation between datasets, respectively (Langfelder *et al*. 2011). Modules were ranked according to their relative preservation using the median rank statistic, which is based on the Z_summary_ score and module size (Langfelder *et al*. 2011).

In addition, we used the NetRep R package (Ritchie *et al*. 2016) to test the significance of all seven of Langfelder’s statistics summarising the preservation of modules between test and discovery datasets. If one or more of the NetRep statistics was found to be non-significant, this was considered evidence for a lack of significant module preservation in subfertile hybrids. The Z_summary_ scores and least significant NetRep statistics for each module are presented in Supplementary Table 1.

### Differential correlation analysis

Genes showing significantly different patterns of pairwise coexpression in the subfertile relative to the fertile hybrids were identified using the R package DGCA (McKenzie *et al*. 2016). Once again, differential correlation analyses were performed independently for each of the F_2_ and HZ populations, and for each of the SFNE and SFAE hybrid groups. The median log-fold change in pairwise coexpression was estimated for each gene in each of the 15 modules, and the significance of median log-fold change values was estimated using 100 permutations.

### Identification of module hub genes

Two methods were used to identify hub genes. First, genes with a Module Membership (kME) ≥ 0.85 were identified as hub genes, where kME represents the Pearson correlation between the expression of an individual gene and the module eigengene (Horvath and Dong, 2008). Second, connectivity statistics including the average number of neighbours, which describes the average connectivity of nodes in a module, and the network density, which summarises the overall module connectivity, were calculated independently for fertile F_2_ and HZ hybrids, using Cytoscape v3.7.1 (Shannon *et al*. 2003). The top five most connected genes within each of the F_2_ and HZ networks were also classified as hub genes for each module.

## Acknowledgments

This work was supported by funding from the Deutsche Forschungsgemeinshaft, awarded to LMT (TU500/2-1). We are grateful to staff at the Milner Centre for Evolution, University of Bath, for valuable feedback and discussions. Expression data processed in this study has been submitted to the Gene Expression Omnibus (GSE136886).

