## Supplementary Figure for "Disrupted gene networks in subfertile hybrid house mice"

Supplementary Table 1. The percentage of the total genes within each module that show significantly different coexpression patterns in subfertile relative to fertile hybrids. Subfertile hybrids are categorized according to the mapping population and the overall expression pattern.

| Module | Percentage genes showing significantly differential correlation | | | | |
|---|---|---|---|---|---|
| | F2:SFAE | F2:SFNE | HZ:SFAE | HZ:SFNE | At least one subfertile group |
| Black | 6.26 | 6.82 | 0.42 | 0.00 | 11.82 |
| Blue | 13.86 | 14.42 | 4.01 | 0.64 | 26.32 |
| Brown | 44.51 | 2.35 | 1.65 | 0.24 | 45.76 |
| Cyan | 2.94 | 7.84 | 0.00 | 2.94 | 10.78 |
| Green | 7.16 | 33.45 | 1.90 | 9.62 | 43.29 |
| Greenyellow | 6.15 | 41.10 | 5.05 | 1.76 | 46.59 |
| Magenta | 2.49 | 1.00 | 0.66 | 0.00 | 3.65 |
| Midnightblue | 8.89 | 22.22 | 2.22 | 0.00 | 28.89 |
| Pink | 15.59 | 3.50 | 9.46 | 2.63 | 25.74 |
| Purple | 0.23 | 1.15 | 0.23 | 0.23 | 1.84 |
| Red | 0.00 | 0.84 | 0.00 | 0.42 | 1.12 |
| Salmon | 2.96 | 4.93 | 4.43 | 0.99 | 11.82 |
| Tan | 14.38 | 7.19 | 4.38 | 5.00 | 25.00 |
| Turquoise | 1.88 | 29.74 | 0.35 | 0.58 | 31.20 |
| Yellow | 3.32 | 22.83 | 8.90 | 16.83 | 41.26 |
| Total Network | 11.08 | 14.83 | 2.88 | 3.06 | 27.53 |

**Supplementary Figure 1.** Principal components analysis (PCA) of genome-wide testis expression data from (A) $F_2$ hybrids and (B) HZ hybrids, performed separately prior to normalization of combined data set and removal of batch effects. Point color indicates fertility class of hybrids, based on sperm count and relative testis weight. (C) Correlation between PC1 loading values in F2 hybrids and HZ hybrids, for 25,146 probes expressed in both datasets.

**Supplementary Figure 2.** Principal components analysis (PCA) of genome-wide expression in testis of pure *Mus musculus domesticus*, *M. m. musculus* and hybrids, PC1 vs. PC3. As indicated in legend, point shape indicates the subspecies or hybrid mapping population for each individual and point colour indicates the fertility class of each hybrid male (see Methods). As indicated in Figure 1, fertile hybrids cluster together with pure subspecies males along the PC1 axis, while males with subfertile phenotypes show higher variability along PC1. PC3 corresponds with subspecies background; *musculus* individuals have high PC3 scores, *domesticus* individuals have low PC3 scores, and hybrids have intermediate scores.

**Supplementary Figure 3.** Moderate disruption of interactions in the Blue module in subfertile $F_2$ hybrids. Although module preservation statistics indicate the Blue module is significantly preserved in SFAE $F_2$ hybrids, several hub genes show loss or weakening of positive correlations. (A) Interactions between module hub genes (red notes) and genes with functions potentially related to male fertility (orange nodes). Gene interactions with an edge-weight exceeding 0.1, as estimated using topological overlap matrices, are indicated using continuous and dashed lines for the fertile and SFAE hybrids, respectively. (B) and (C) Coexpression heatmaps showing pairwise correlations between expression values of Blue module genes in fertile and SFAE $F_2$ hybrids, respectively.

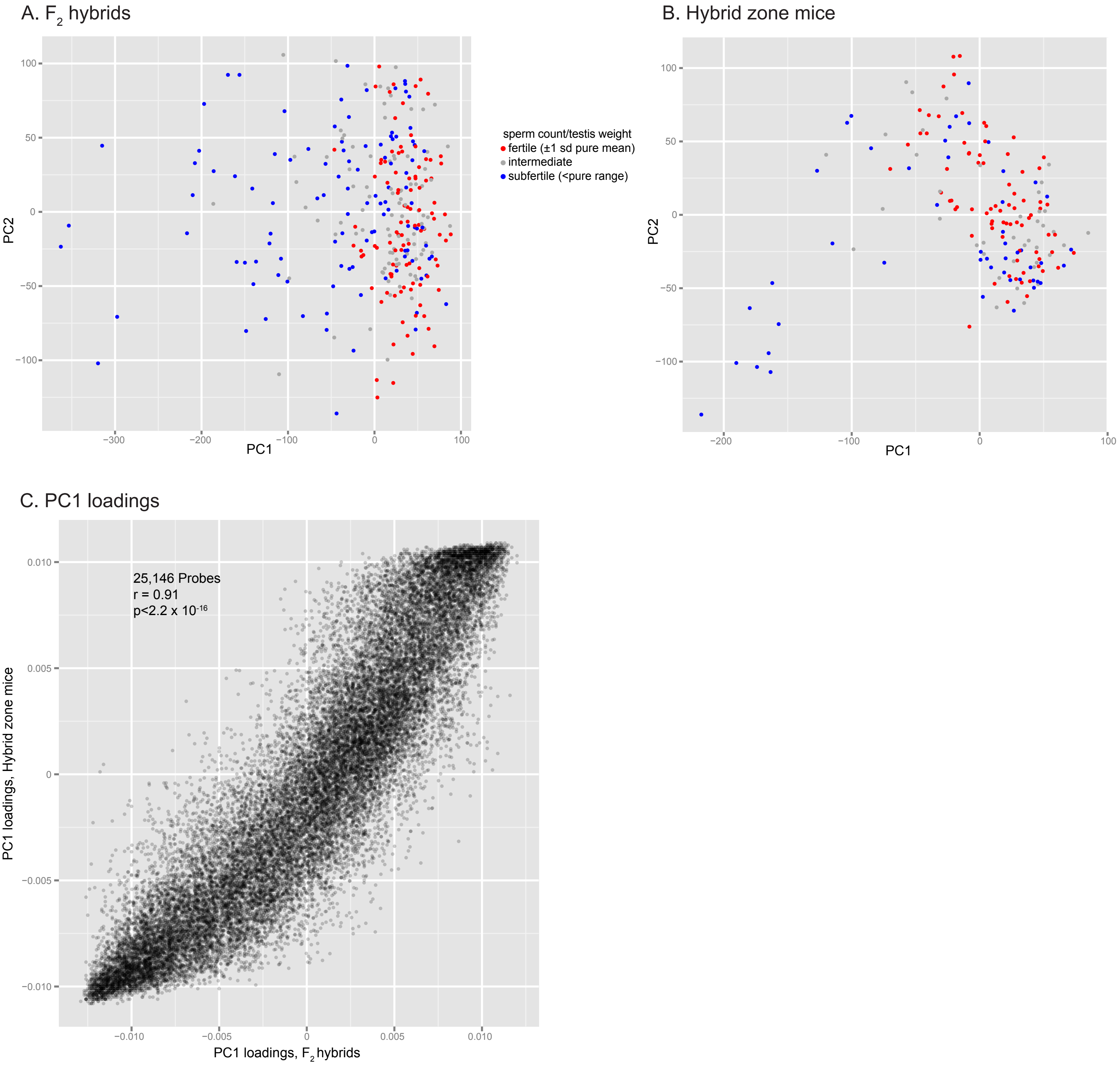
A. F2 hybrids
PC1
PC2
B. Hybrid zone mice
PC1
PC2
sperm count/testis weight
fertile (±1 sd pure mean)
intermediate
subfertile (<pure range)
C. PC1 loadings
25,146 Probes
r = 0.91
p<2.2 x 10-16
PC1 loadings, F2 hybrids
PC1 loadings, Hybrid zone mice

A.

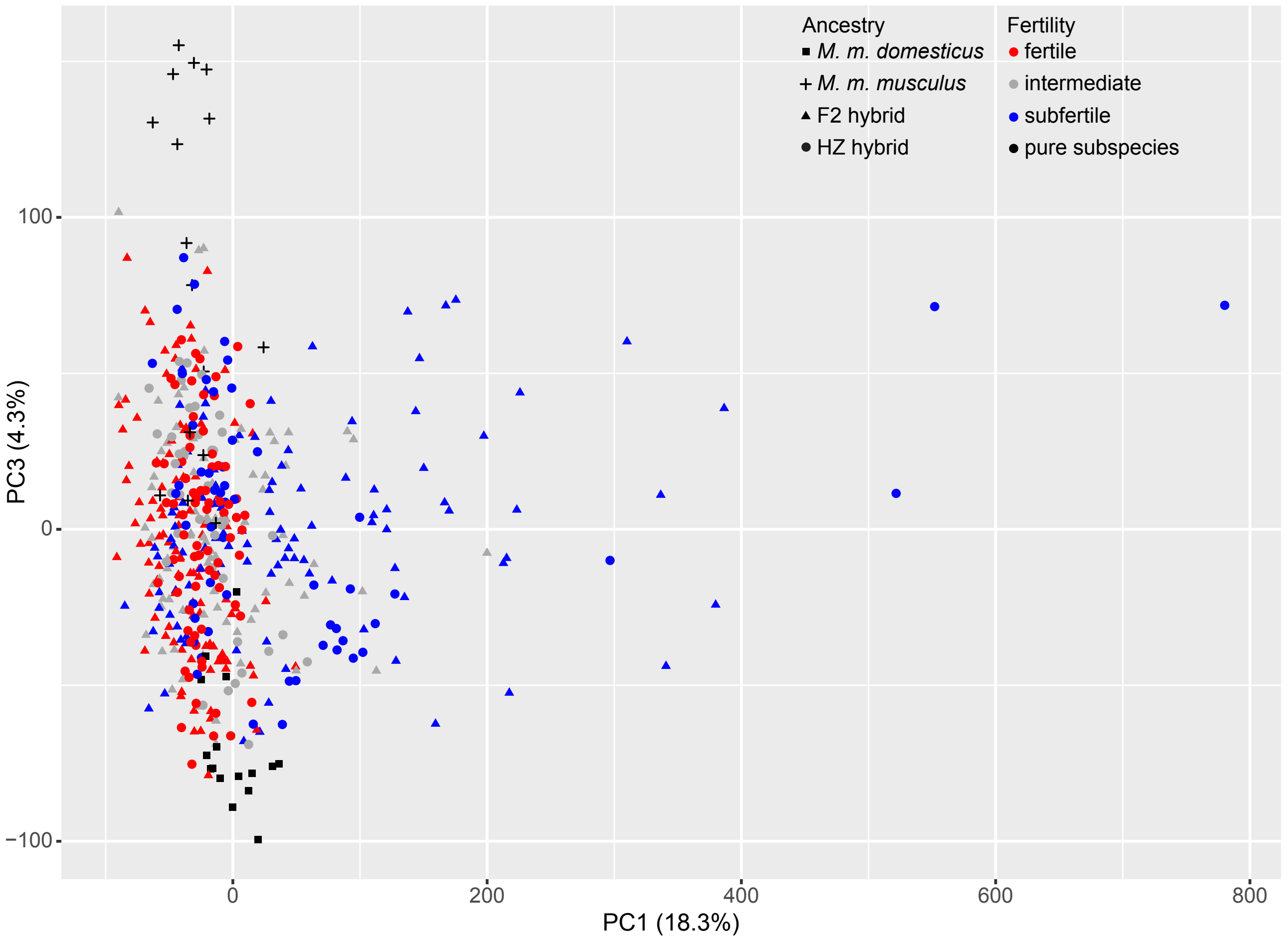
Ancestry
M. m. domesticus
M. m. musculus
F2 hybrid
HZ hybrid
Fertility
fertile
intermediate
subfertile
pure subspecies
PC3 (4.3%)
100
0
−100
0
200
400
600
800
PC1 (18.3%)

B.

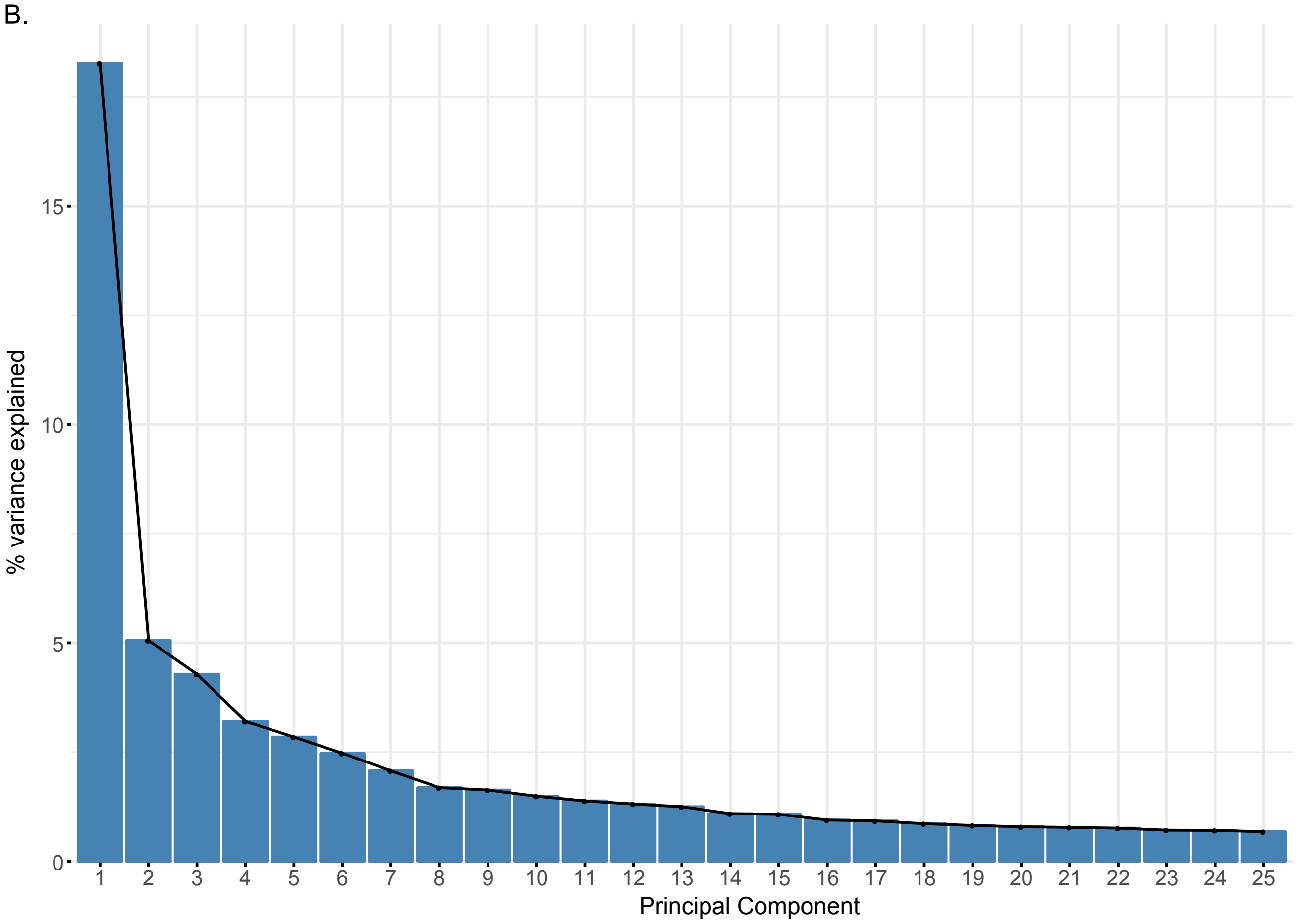
% variance explained
15
10
5
Principal Component

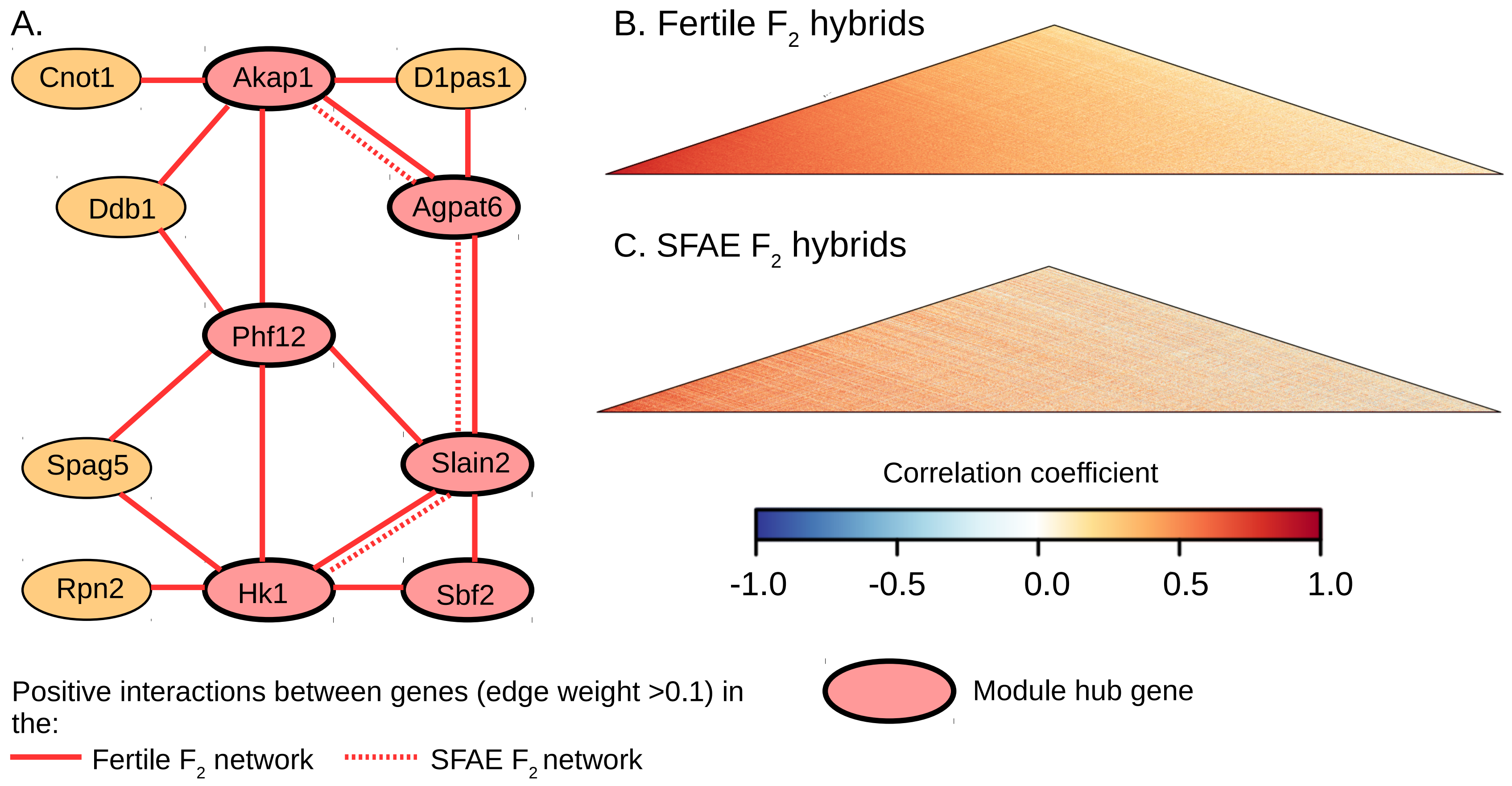
A.
Cnot1
Akap1
D1pas1
Ddb1
Agpat6
Phf12
Spag5
Slain2
Rpn2
Hk1
Sbf2
B. Fertile $F_2$ hybrids
C. SFAE $F_2$ hybrids
Correlation coefficient
-1.0
-0.5
0.0
0.5
1.0
Positive interactions between genes (edge weight >0.1) in the:
Fertile $F_2$ network
SFAE $F_2$ network
Module hub gene
